# Inhibitory effect of Dolosigranulum pigrum and Corynebacterium pseudodiphtheriticum on pneumococcal in vitro growth

**DOI:** 10.1101/2025.01.10.632320

**Authors:** M Cisneros, M Blanco-Fuertes, A Lluansí, P Brotons, D Henares, A Pérez-Argüello, G González-Comino, P Ciruela, A Mira, C Muñoz-Almagro

## Abstract

**Background:** *Streptococcus pneumoniae* is a nasopharynx coloniser that can invade sterile tissues, causing Invasive Pneumococcal Disease (IPD). *Dolosigranulum pigrum* and *Corynebacterium pseudodiphtheriticum* are commensal bacteria commonly isolated from the nasopharynx of healthy children, potentially playing a protective role. This study aims to analyse the effects of *D. pigrum* and *C. pseudodiphtheriticum* on *S. pneumoniae* in vitro growth.

**Methods:** Pneumococcal strains were collected from IPD patients and healthy carriers in Catalonia (2016-2023). *D. pigrum* and *C. pseudodiphtheriticum* strains were isolated from a healthy child’s nasopharynx. *S. pneumoniae* was co-cultured with each commensal bacterium in triplicate experiments. Pneumococcal growth was quantified using a real-time PCR assay targeting the *lytA* gene. The effect of commensal bacteria on pneumococcal growth was evaluated using a linear mixed-effect regression model.

**Results:** Twenty-eight pneumococcal strains expressing 24 different serotypes and 26 clonal types were analysed (18 isolated in blood and 10 in nasopharyngeal aspirate). Pneumococcal growth was decreased by *D. pigrum* (β = −0.763, 95% CI: −0.94 to −0.59, p < 0.0001) and *C. pseudodiphtheriticum* (β = −0.583, 95% CI: −0.76 to −0.41, p < 0.0001). The combined presence of both had a stronger inhibitory effect (β = −0.971, 95% CI: −1.15 to −0.79, p < 0.0001). No association was found between isolation site or serotype with pneumococcal growth.

**Conclusion:** *D. pigrum* and *C. pseudodiphtheriticum* significantly reduced pneumococcal growth, with a synergic effect when combined. This antagonistic effect supports the potential protective factor of healthy nasopharyngeal microbiota against IPD and the development of these microorganisms as probiotics.

## INTRODUCTION

*Streptococcus pneumoniae* is a gram-positive bacterium that asymptomatically colonises the human nasopharynx, especially among children aged under 5 years. However, under certain circumstances, it has the potential to invade sterile tissues and cause disease (1). Invasive Pneumococcal Disease (IPD) is the most severe form of pneumococcal infection and includes bacteremia, pneumonia, meningitis, and septicemia, among others. Despite the availability of pneumococcal vaccines, it has been estimated that *S. pneumoniae* is responsible for over 300,000 deaths annually in children under five years of age (2).

Pneumococcal conjugate vaccines (PCVs) are developed against the bacterium’s main virulence factor, the capsular polysaccharide (CPS) (3). There are more than 100 known capsular serotypes with different degrees of invasiveness; some of them are high invasive disease potential serotypes (4–6). Following the introduction of conjugate vaccines, numerous studies have shown a decreased prevalence of vaccine serotypes (VTs) and a significant reduction in IPD caused by these serotypes. Nevertheless, vaccines do not cover all serotypes, leading to the phenomenon of serotype replacement, in which non-vaccine serotypes (NVTs) are expanded and become more in disease (7, 8).

Pneumococcal nasopharyngeal colonisation represents a key factor to understand the burden of pneumococcal disease and address its prevention. Colonisation is a complex and dynamic process in which multiple factors interplay, including environmental, host, and microbiota factors. Intricate interactions with other common inhabitants of the nasopharynx seem to be an important step for pneumococcal colonisation (1, 9).

Previous studies have suggested that certain natural colonisers of the nasopharynx could play a protective role against the colonisation of *S. pneumoniae* (10). Significant abundances of *Dolosigranulum pigrum* and *Corynebacterium* species have been found in nasal and nasopharyngeal microbiota of children when *S. pneumoniae* is absent (10–12). Furthermore, increased abundances have been detected in healthy controls compared to patients with different respiratory infections, suggesting a potential beneficial role in respiratory health through negative interactions with common pathobionts (10–19).

*D. pigrum* is one of the known species of the genus *Dolosigranulum* that was first reported (20) (21). In terms of structure, it is a gram-positive coccus arranged in pairs, tetrads, and clusters. This catalase-negative bacterium is generally sensitive to antibiotics like beta-lactams (22). A notable characteristic of *D. pigrum* that enhances it as a potential protective species is its ability to produce lactate, classifying it as lactic acid bacterium (LAB) (23, 24). Additionally, genome sequencing has confirmed this classification by identifying genes associated with the homofermentation of carbohydrates (24, 25). Most LABs have been found in the digestive tract, where they perform beneficial tasks for human health, such as enhancing resistance to pathogens through microbe-microbe interactions, the synthesis of bacteriocins, or immunomodulation (26). For instance, *Lactobacillus murinus* a LAB from the lung microbiota in mice can inhibit *S. pneumonia* growth in vitro and prevent colonization in vivo (27).

*D. pigrum* is commonly isolated from the nasal cavity or the nasopharynx, where it coexists with other commensal bacteria such as *Corynebacterium spp* (11). It has been also described that *Corynebacterium pseudodiphtheriticum*, one of the species of this genus, could also prevent pathogenic bacteria colonisation in the nasopharynx in cooperation with *D. pigrum* (28). Brugger S, et al. demonstrated increased relative abundances of *C*. *pseudodiphtheriticum* were increased in the presence of *D. pigrum*. Additionally, they observed that *C*. *pseudodiphtheriticum* could enhance the growth of *D. pigrum* in vitro (14).

Despite these suggestive results, little is known about the impact of *D. pigrum* and *C. pseudodiphtheriticum* on pneumococcal replication rate and the consistency of these results according to different capsular serotypes of *S. pneumoniae*. Thus, the aim of the present study is to analyse the effects of the two commensal bacteria, *D. pigrum* and *C. pseudodiphtheriticum* on *Streptococcus pneumoniae* in vitro growth (IVG) in a diverse collection of *S. pneumoniae* strains.

## MATERIAL AND METHODS

### Bacterial strains collections

Invasive pneumococcal strains were obtained from the collection of the Support Laboratory for Molecular Epidemiology of IPD in Catalonia during 2016-2023. Carriage strains (*D. pigrum, C. pseudodiphtheriticum, and S. pneumoniae*) were prospectively collected with informed consent in the framework of two consecutive funded projects focused on the role of nasopharyngeal microbiota in respiratory health and disease during 2016-2023. All the strains were preserved in 1 ml of preservation media of skim milk at −80°C.The selected strains represented the main serotypes and clones detected in Catalonia during the study period as reported on Microbiological notification system of Catalonia.

### Isolation and identification process of *Dolosigranulum pigrum* and Corynebacterium pseudodiphtheriticum

*D. pigrum* and *C. pseudodiphtheriticum* strains were isolated from a nasopharyngeal aspirate (NPA) sample of a healthy child attended at Sant Joan de Déu Barcelona Children’s Hospital (SJD) in the autumn of 2018. Isolation of both bacteria was performed by culturing 100 µl of NPA sample on Blood Agar plates (Columbia agar supplemented with 5% sheep blood; BioMérieux). Additionally, *D. pigrum* was isolated on CNA Agar plates (Columbia CNA agar supplemented with 5% sheep blood; Becton Dickinson) and Mannitol Salt Agar (Becton Dickinson). The plates were incubated at 37 °C under aerobic conditions supplemented with 5% CO^2^.

MALDI-TOF mass spectrometry was used to identify both commensal bacteria. This system generates a spectrum based on the mass-charge relationship of the microorganism’s proteins, which is then compared with reference libraries containing the different spectra of known microorganisms (29).

### Isolation, identification, and molecular characterisation process of Streptococcus pneumoniae

Pneumococcal isolates were obtained from blood and nasopharyngeal samples from IPD patients and healthy carriers in Catalonia during 2016–2023.

To isolate *S. pneumoniae,* 100 µl of the NPAs were directly cultured in Blood Agar plates (Columbia agar supplemented with 5% sheep blood; BioMérieux) and incubated at 37 °C under aerobic conditions supplemented with 5% CO^2^. Pneumococcal identification was conducted using standard microbiological techniques, such as the optochin sensitivity test and colony morphology (30). In addition, capsular typing was performed on all pneumococcal using fluorescence fragment analysis and Whole Genome Sequencing (WGS) (31, 32). According to previous literature, (5) Serotypes 1, 3, 4, 5, 7F, 8, 9A, 9V, 12F, 14, 18, 19A, 24F, and 33F were considered high invasive disease potential serotypes (HIPST) while the rest were considered low invasive disease potential serotypes (LIPST).

### In vitro growth of pneumococcal isolates according to *D. pigrum* and *C. pseudodiphtheriticum* exposure

#### Preparation and standardisation of D. pigrum and C. pseudodiphtheriticum enriched media

Isolates of *D. pigrum* and *C. pseudodiphtheriticum* preserved at −80°C were cultured on Blood Agar plates (Columbia agar supplemented with 5% sheep blood; BioMérieux). To establish different in vitro conditions for *S. pneumoniae*, overnight cultures of *D. pigrum* and *C. pseudodiphtheriticum* (grown in aerobic conditions with 5% CO^2^ at 37 °C) were suspended in 2 ml of fresh BBL Todd-Hewitt broth (Becton Dickinson). The suspension solution was adjusted to a concentration of 6 log_10_ genome copies per microlitre ± 0.7 log_10_, determined by Qubit Fluorometric Quantification from the extraction of genomic DNA ***(Figure 1)*.**

**Figure 1:**
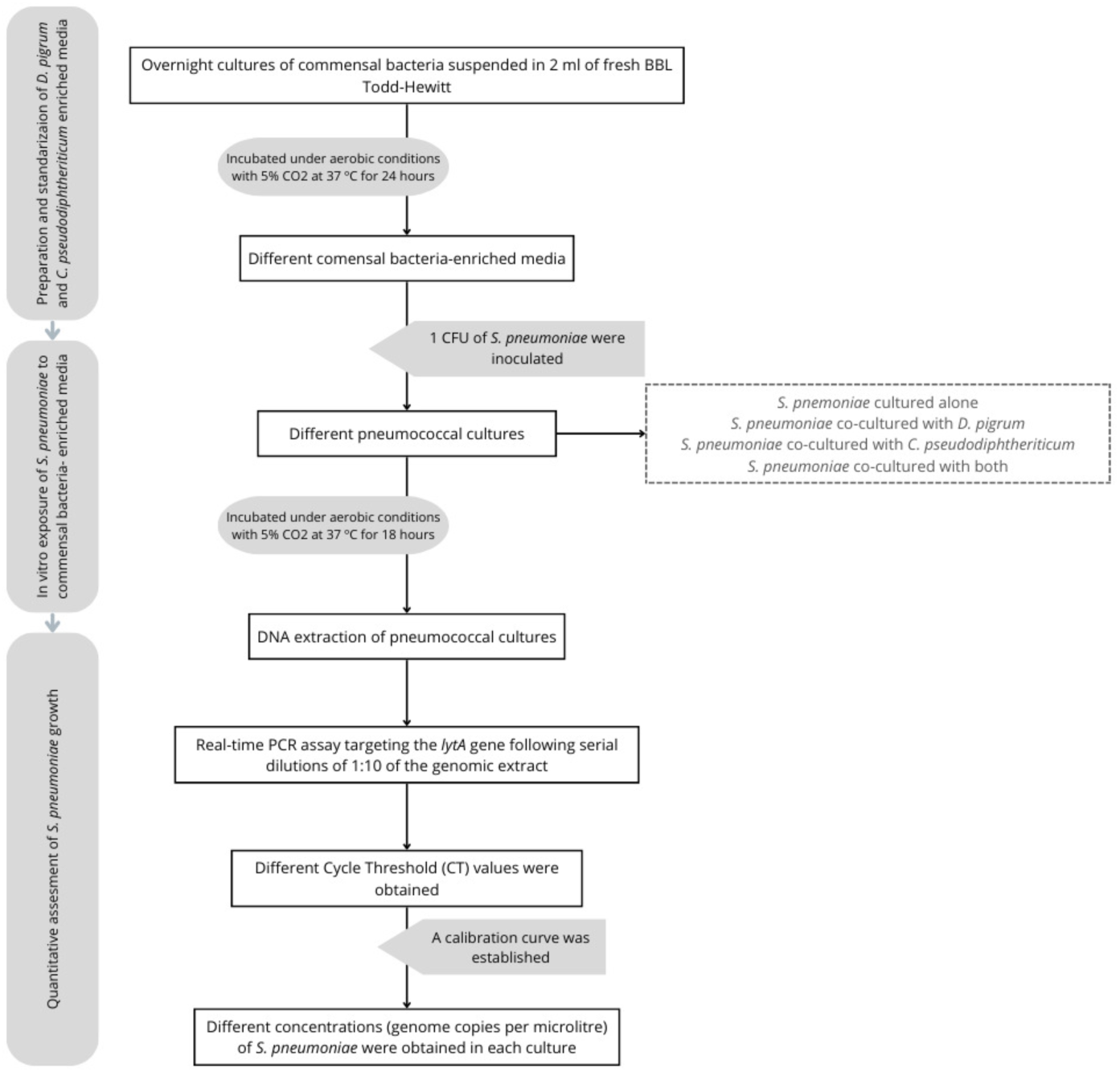
Flow diagram of the protocol for in vitro growth of pneumococcal isolates according to *D. pigrum* and *C. pseudodiphtheriticum* exposure.

#### In Vitro Exposure of S. pneumoniae to Commensal Bacteria-Enriched Media

In order to expose *S. pneumoniae* to commensal bacteria, 1 CFU of each pneumococcal isolate was inoculated into four culture conditions, resulting in the following pneumococcal broth cultures: (i) *S. pneumoniae* cultured alone in 2 ml of BBL Todd-Hewitt broth (Becton Dickinson) as negative controls (S); (ii) *S. pneumoniae* co-cultured with *D. pigrum* (SD); (iii) *S. pneumoniae* co-cultured with *C. pseudodiphtheriticum* (SC); and (iv) *S. pneumoniae* co-cultured with both (SDC). Cultures were incubated under aerobic conditions with 5% CO^2^ at 37 °C for 18 hours. To account for the intrinsic heterogeneity among strains, each strain was tested in triplicate ***(Figure 1)*.**

#### Quantitative Assessment of S. pneumoniae Growth

DNA was extracted from 400 µl of each of the four pneumococcal cultures using the eMAG platform (bioMerieux; Marcy-l’Étoile, France) with a final elution volume of 50 µl.

*S. pneumoniae* was quantified using a real-time PCR assay targeting the *lytA* gene following serial dilutions of 1:10 of the genomic extract. Primers and probes were employed according to the Centers for Disease Control and Prevention (CDC) guidelines (33).

A calibration curve was established to correlate *S. pneumoniae* DNA concentration (ng/µl) with the Cycle Threshold (CT) values obtained from the real-time PCR. The calibration curves were generated from the extraction of genomic DNA from a Todd-Hewitt suspension of *S. pneumoniae* of each strain. The DNA concentration was measured using Qubit Fluorometric Quantification with serial 1:10 dilutions prepared from the range of 10 to 10^-6^ nanograms per microliter (ng/µl). The resulting pneumococcal IVG values, initially quantified in ng/ µl, were converted to gc/µl using the median genome size of *S. pneumoniae*, which is 2.085 kbp (34) ***(Figure 1)*.**

### Statistical analyses

The effect of commensal bacteria on pneumococcal growth was considered according to microbial and clinical characteristics of pneumococcal infection: serotype invasiveness, healthy pneumococcal nasopharyngeal carriage or IPD status, and isolation site (blood or nasopharynx).

A linear mixed-effect regression model (LMM) was used to evaluate the impact of *D. pigrum* and *C. pseudodiphtheriticum* on *S. pneumoniae*’s growth. To normalise the distribution, the outcome variable, pneumococcal growth, underwent a log transformation. The strain identifier and the strain replicate were used as nested random effects. This approach allows for control and measurement of the inherent differences in pneumococcal growth between strains and ensures not biased results derived from fixed effects.

All non-collinear variables examined—the culture condition, serotype invasiveness, and site of isolation—were included in an initial LMM. To determine which variables should be used as a fixed effect, a stepwise approach from the initial LMM was used to iteratively remove variables, resulting in different models. The Akaike Information Criteria (AIC), Restricted Maximum Likelihood (REML), and Determination Coefficient (R²) (35–37) were the three key adjustment criteria used in each step to assess model fit and identify the significant predictor variables. This process led to the selection of the optimal LMM, the LMM2.

To assess the collinearity among the fixed effect variables, the Variance Inflation Factor (VIF) was used, and those with a VIF > 5 were deemed collinear (38). Associations between these variables were measured by Estimated coefficient (β), and their significance was assessed by estimated Marginal Means. 95% confidence intervals (CIs) were calculated, and statistical significance was determined by setting a p-value of 0.05.

All the statistical analyses were performed using the 4.3.1 version of R and RStudio software (39) using the *car* (40), *lme4* (41), *emmeans* (42), *lmerTest* (43), and *MumIn* (44) packages.

## RESULTS

### Pneumococcal strain’s features

During the study period (2016-2023), a total of 28 *S. pneumoniae* strains were selected for analysis; 18 were isolated from blood and 10 from nasopharyngeal samples (2 from IPD patients and 8 from healthy children). Twenty-four different serotypes and 27 different clones were detected in the 28 pneumococcal strains. A proportion of 60.71% of serotypes (n=17) were classified as low invasive disease potential serotypes ***(Table 1)*.**

**Table 1:** Characteristics of Pneumococcal strains included in this study.

| Strain | Serotype | Invasiveness of<br>pneumococcal<br>serotypes | Clone | Year of<br>isolation | Site of<br>isolation | Health<br>status |
| --- | --- | --- | --- | --- | --- | --- |
| 1 | 10A | LIPST | 97 | 2015 | NPA | Healthy |
| 2 | 9N | LIPST | 66 | 2016 | NPA | IPD |
| 3 | 6C | LIPST | 4310 | 2016 | NPA | Healthy |
| 4 | 6B | LIPST | 13321 | 2016 | NPA | Healthy |
| 5 | 15B | LIPST | 1262 | 2017 | NPA | Healthy |
| 6 | 20 | LIPST | 1871 | 2017 | NPA | Healthy |
| 7 | 19F | LIPST | 1167 | 2017 | NPA | Healthy |
| 8 | 15A | LIPST | 5139 | 2018 | NPA | Healthy |
| 9 | 33F | HIPST | 717 | 2018 | NPA | Healthy |
| 10 | 11A | LIPST | 6521 | 2018 | NPA | IPD |
| 11 | 5 | HIPST | 289 | 2017 | Blood | IPD |
| 12 | 2 | LIPST | 74 | 2019 | Blood | IPD |
| 13 | 6A | LIPST | NA | 2019 | Blood | IPD |
| 14 | 23F | HIPST | 277 | 2021 | Blood | IPD |
| 15 | 7F | HIPST | 3544 | 2021 | Blood | IPD |
| 16 | 1 | HIPST | 306 | 2023 | Blood | IPD |
| 17 | 22F | LIPST | 433 | 2022 | Blood | IPD |
| 18 | 18C | HIPST | 133 | 2023 | Blood | IPD |
| 19 | 14 | HIPST | 156 | 2023 | Blood | IPD |
| 20 | 19A | HIPST | 695 | 2023 | Blood | IPD |
| 21 | 4 | HIPST | 205 | 2023 | Blood | IPD |
| 22 | 11A | LIPST | 6521 | 2023 | Blood | IPD |
| 23 | 8 | HIPST | 53 | 2023 | Blood | IPD |
| 24 | 12F | HIPST | 3377 | 2023 | Blood | IPD |
| 25 | 9N | LIPST | 517 | 2023 | Blood | IPD |
| 26 | 3 | HIPST | 180 | 2023 | Blood | IPD |
| 27 | 17F | LIPST | 392 | 2023 | Blood | IPD |
| 28 | 20 | LIPST | 15084 | 2023 | Blood | IPD |

NA: Not available; IPD: Invasive Pneumococcal Disease; NPA: Nasopharyngeal aspirate; HIPST: High invasive disease potential serotypes; LIPST: Low invasive disease potential serotypes

### Descriptive results

All 28 pneumococcal strains were replicated three times, resulting in a total of 84 observations. Mean growth of pneumococcal isolates in monoculture (3.04 log_10_gc/µl; SD, 0.42) was higher compared to those co-cultured with *D. pigrum* (2.28 log_10_gc/µl; SD, 0.57), *C. pseudodiphtheriticum* (2.46 log_10_gc/µl; SD, 0.82), or both commensal species (2.07 log_10_cg/µl; SD, 0.78) ***(Table 2).*** The mean growth of high invasive disease potential serotypes (2.61 log_10_gc/µl; SD, 0.75) pneumococcal isolates was higher than low invasive disease potential serotypes (2.35 log_10_gc/µl; SD, 0.74). Similarly, pneumococcal isolates from blood samples showed a higher mean growth (2.53 log_10_gc/µl; SD, 0.72) compared to those from NPA (2.32 log_10_gc/µl; SD, 0.79) ***(Table 2)***.

**Table 2:** Mean bacterial culture growth of 28 pneumococcal strains according to different categorical variables under various culture conditions.

| Pneumococcal growth (log<sub>10</sub> gc/μl) (Mean+ sd) |  |  |  |  |  |
| --- | --- | --- | --- | --- | --- |
|  | *S. pneumoniae*<br>(S) | *S. pneumoniae*<br>+ *D. pigrum*<br>(SD) | *S. pneumoniae* +<br>*C. pseudodiphtheriticum*<br>(SC) | *S. pneumoniae* + *D. pigrum*<br>+<br>*C. pseudodiphtheriticum*<br>(SDC) | Overall |
| Overall | 3.041 (0.422) | 2.279 (0.569) | 2.459 (0.815) | 2.071 (0.775) |  |
| HIPST | 3.106 (0.453) | 2.447 (0.533) | 2.614 (0.920) | 2.287 (0.787) | 2.613 (0.757) |
| LIPST | 2.999 (0.400) | 2.171 (0.570) | 2.359 (0.732) | 1.933 (0.742) | 2.365 (0.738) |
| NPA | 2.986 (0.475) | 2.127 (0.624) | 2.191 (0.884) | 2.094 (0.762) | 2.350 (0.787) |
| Blood | 3.071 (0.391) | 2.363 (0.523) | 2.608 (0.742) | 2.059 (0.789) | 2.525 (0.729) |

sd: standard deviation

### Selection of optimal model

Three LMMs were conducted to identify which predictor variables of *S. pneumoniae* strains had an effect on their growth. LMM2 presented lower AIC (580.748) and REML (564.7) values, indicating better fit compared to LMM1 and LMM3. Although LMM2 did not present the greatest Marginal R^2^ value, the difference from the others is minimal. All the models had similar conditional values of Conditional R², suggesting that the overall explanatory power of the models remains unaffected by the addition of new predictor variables. For this reason, LMM2 provides a good balance between model fit and the inclusion of relevant predictors ***(Table 3 and S1)***.

**Table 3:** The effect of commensal bacteria to pneumococcal growth of 28 strains of *S. pneumoniae* using linear mixed-effect model.

| Predictors | $\beta$ | 95% CI | SE | p value |
| --- | --- | --- | --- | --- |
| Intercept | 3.195 | 2.91, 3.479 | 0.146 | $p < 0.0001$ **** |
| Culture condition |  |  |  |  |
| SD vs S | -0.763 | -0.941, -0.586 | 0.069 | $p < 0.0001$ **** |
| SC vs S | -0.583 | -0.761, -0.406 | 0.069 | $p < 0.0001$ **** |
| SDC vs S | -0.971 | -1.148, -0.793 | 0.069 | $p < 0.0001$ **** |
| SD vs SC | -0.180 | -0.357, -0.003 | 0.069 | 0.045 * |
| SDC vs SD | -0.207 | -0.385, -0.030 | 0.069 | 0.0146 * |
| SDC vs SC | -0.387 | -0.565, -0.210 | 0.069 | $p < 0.0001$ **** |
| Invasiveness of serotype<br>(LIPST vs HIPST) | -0.250 | -0.601, 0.101 | 0.180 | 0.175 |
| **Random Effects** |  |  |  |  |
| $\sigma^2$ | 0.197 | | | |
| $\tau_{00}$ Replicate:Strain_ID | 0.063 | | | |
| $\tau_{00}$ Strain_ID | 0.178 | | | |
| ICC | 0.550 |  |  |  |
| $N_{\text{REPLICATE}}$ | 3 | | | |
| $N_{\text{STRAIN}}$ | 28 | | | |
| Observations | 336 |  |  |  |

$\beta$ : Estimated Coefficient; SE: Standard Error; $\sigma^2$ : Residual variance; $\tau_{00}$ : Intercept variance; ICC: Intraclass Correlation Coefficient

### Pneumococcal strain variability

The results of the LMM2 revealed considerable variability in pneumococcal growth among strains. The intraclass correlation coefficient (ICC) was 0.55, indicating that 55% of the total variability in growth could be attributable to differences between individual strains. The variance of the random effects between strains was 0.178, while the variance between replicates within the same strain was lower, 0.063 ***(Table 3).*** These findings showed that a considerable portion of the observed variability in pneumococcal growth was accounted for by differences between individual strains rather than variation among replicates of the same strain.

### Inhibitory effect of Dolosigranulum pigrum and Corynebacterium pseudodiphtheriticum on pneumococcal growth

The results of LMM2 analysis provided strong evidence of the association between the culture condition and the pneumococcal growth, although pneumococcal strain variability was observed.

The presence of *D. pigrum* was significantly associated with a reduction in pneumococcal growth, as reflected by the estimate coefficient (β = −0.763, CI: −0.94 to −0.59, p < 0.0001). Similarity *C. pseudodiphtheriticum* exhibited a notable inhibitory effect, (β = −0.583, CI: −0.76 to −0.41, p < 0.0001). Notably, the combined presence of both bacteria was strongly associated (β = −0.971, CI: −1.15 to −0.79, p < 0.0001), with pneumococcal growth decrease.

Even though *D. pigrum* and *C. pseudodiphtheriticum* showed an individual inhibitory effect on pneumococcal growth, their combined effect was even significantly stronger, with estimated coefficients of −0.207 (CI: −0.39 to −0.03, p < 0.01) and −0.387 (CI: −0.57 to −0.21, p < 0.0001) when compared to co-culture with *D. pigrum* alone or *C. pseudodiphtheriticum* alone, respectively ***(Figure 2 and Table 3)***.

**Figure 2:**
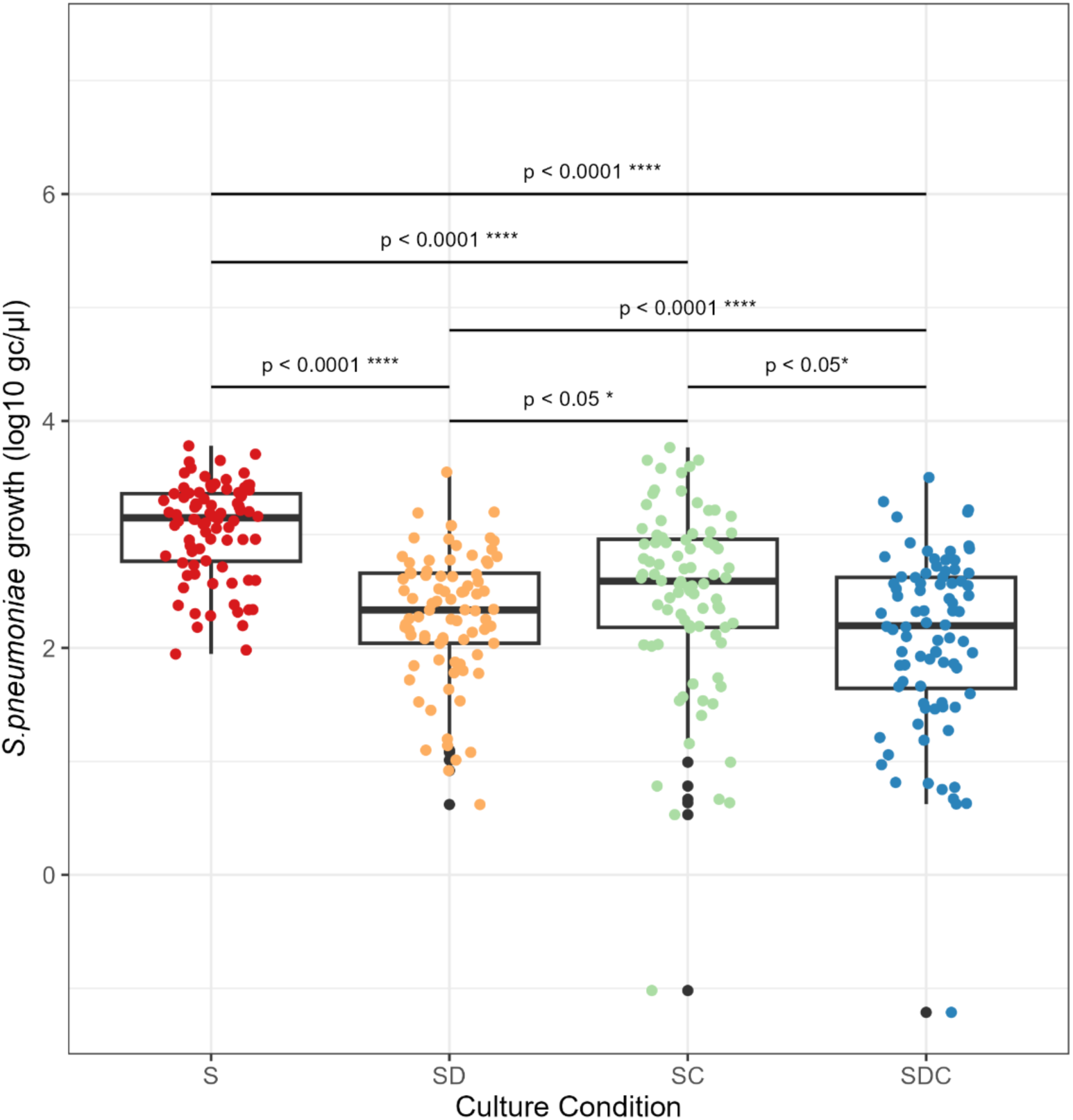
Pneumococcal growth under different culture conditions: Cultured alone (red, labeled as S for *Streptococcus*), cultured with *D. pigrum* (orange, labeled as SD), cultured with *C. pseudodiphtheriticum* (light green, labeled as SC), and cultured with both commensal bacteria (dark blue, labeled as SDC). The data is represented by boxplots showing the median and interquartile range (IQR) of pneumococcal growth (measured in gc/¿l after log-transformation), along with individual data points (n=338). Black points denote outliers. *\*p < 0.05; **p < 0.01; ***p < 0.001; ****p < 0.0001*.

### Variation in pneumococcal growth according to strain features

In the LMM analysis, the site of isolation did not indicate any significant association with pneumococcal growth (β = −0.079, CI = −0.48 to 0.33, p = 0.708) ***(Figure 3C and Table S1)***. Furthermore, pneumococcal growth appeared not to be generally affected by the invasiveness of pneumococcal serotype. low invasive disease potential serotypes pneumococcal isolates showed a slightly reduced growth, although not statistically significant, overall compared to high invasive disease potential serotypes (β = −0.250, CI = −0.60 to −0.10, p = 0.175) ***(Figure 3A and Table 3)***. Moreover, a detailed subanalysis examining this predictor under different culture conditions revealed that low invasive disease potential serotypes isolates showed a greater decrease in growth when they were cultured with both commensal bacteria (β = −0.355, CI = −0.76 to 0.05, p = 0.082) than when they were cultured alone, where the effect was minimal (β = −0.112, CI = −0.52 to 0.29, p = 0.577). However, this effect was not significant, indicating no clear association between these variables. ***(Figure 3B and Table S2)***.

**Figure 3:**
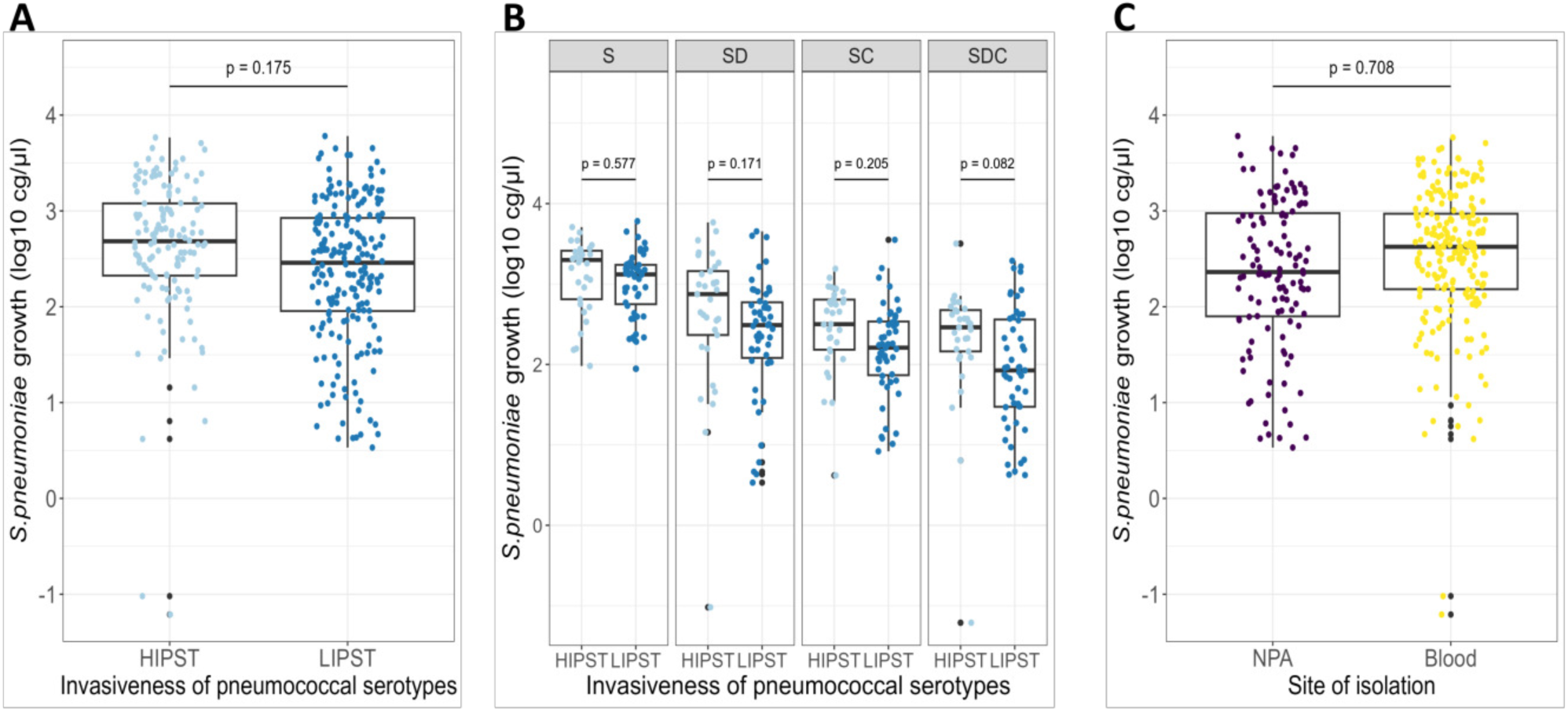
Pneumococcal growth according to strain features. **A)** Invasiveness of pneumococcal serotypes, HIPST (light blue) vs LIPST (dark blue). **B)** Invasiveness of pneumococcal serotypes in different culture conditions. **C)** Site of isolation, NPA (purple) vs blood (yellow). The data is represented by boxplots showing the median and interquartile range (IQR) of pneumococcal growth (measured in gc/¿L after log-transformation), along with individual data points (n=338). Black points denote outliers. *\*p < 0.05; ** p < 0.01; *** p < 0.001; **** p < 0.0001*

## DISCUSSION

The present study evaluated the effect of two commensal bacteria from the nasopharyngeal microbiota on the growth of different pneumococcal strains as well as other characteristics of the strain itself, such as serotype invasiveness and the isolation site of the isolate, which may influence this effect.

The main findings of this study indicated that despite the variability among pneumococcal strains, the presence of those two commensal bacteria, alone or in combination, had a substantial impact on pneumococcal growth. Specifically, the growth of *S. pneumoniae* was significantly decreased in the presence of *D. pigrum,* supporting its potential interest as a candidate for inclusion in a probiotic formula. An ideal probiotic for the respiratory microbiota should be a non-pathogenic bacterium that is present as a natural carrier, capable of adhering to the epithelium and colonising the niche of key pathobionts. It also should have no cytotoxic effect on respiratory epithelial cells, resistance against the horizontal gene transfer and mobile genetic elements, a low tendency for tissue invasion, and be susceptible to commonly used antibiotics (19, 26). Recent reviews of the literature highlight these characteristics of *D. pigrum*, supporting its potential as a probiotic candidate (19).

Nonetheless, even with these promising features, the current published studies have been confined to very few strains and have not considered the extensive genetic diversity of *S. pneumoniae,* which consists of 100 serotypes and thousands of clones with different invasive disease potential (1, 2, 4, 45).

One of the key strengths of this study is the use of various strains of *S. pneumoniae* with different intrinsic characteristics, such as distinct serotypes, and isolated from different anatomical sites. This diversity in pneumococcal strains allowed considering strain-specific differences in the pneumococcal growth in the analysis, providing stronger insights into potential interactions.

The analysis conducted on *S. pneumoniae* strain-specific variables revealed that the site of isolation (nasopharynx or blood) and serotype invasiveness did not significantly affect the antagonistic effect of commensal bacteria in pneumococcal growth. These findings suggest that the inhibition of *S. pneumoniae* is consistent regardless of its site of isolation and invasiveness, enhancing the capacity of *D. pigrum* as a potential probiotic. However, the limited number of strains emphasises the importance of additional studies to validate this consistent antagonistic effect and thus better comprehend the underlying interactions.

The exact mechanisms by which *D. pigrum* exerts its observed antagonistic effect on pneumococcal growth remain unclear. One possible explanation is competition of nutrients, which are frequently a limiting factor for bacteria colonisation, such as *S. pneumoniae.* Therefore, *S. pneumoniae* experiences reduced nutritional availability when it coexists with other bacteria, which consequently restricts its growth (46).

Another factor could be the secretion of antimicrobial compounds, such as bacteriocins, which can function as a bactericidal or bacteriostatic agent against pathogens (23, 24). Specifically, it is suggested that *D. pigrum* may produce lantipeptides, a type of bacteriocin with notable antimicrobial activity that can disrupt bacteria’s cell walls, restricting pathogen proliferation (47–49).Thus, pneumococcal inhibition may be explained in part by *D. pigrum’s* antimicrobial characteristics. In order to comprehend the precise association between *D. pigrum* and *S. pneumoniae* and the role of bacteriocins in this instance, additional analyses are crucial.

*D. pigrum* as a LAB, has the ability to generate lactate in the nasal microbiome. Lactate is an acid that can lower the environmental pH, creating unfavourable conditions for the optimal growth of *S. pneumoniae*, which grows in a more neutral pH (50). However, prior studies such as those by Brugger S, et al., indicate that *D. pigrum*’s lactate production for itself is insufficient to fully inhibit pneumococcal growth (14). This suggests that additional inhibition mechanisms are taking place.

An interesting aspect of the results is the synergistic effect observed when another commensal bacterium was present. Specifically, the inhibition of *S. pneumoniae* growth was significantly greater when *D. pigrum* cooperated with *C. pseudodiphtheriticum*. Although both bacteria individually already showed inhibitory effects, their combined effect is significantly stronger. This suggests that the interaction between *D. pigrum* and *C. pseudodiphtheriticum* enhances the reduction of pneumococcal growth. This observation could be indicative of a synergistic interaction between these two commensal bacteria, which aligns with the study of Brugger S. In this regard, *D. pigrum* is auxotrophic for certain nutrients, especially for aminoacids, and it is hypothesised that *C. pseudodiphtheriticum* may provide these nutrients, enhancing their mutual inhibitory effects on *S. pneumoniae* (14, 51, 52). A synergistic effect of bacteriocins from different bacterial isolates cannot be ruled out either.

Some limitations should be noted in the study. First, only a single strain of *D. pigrum* and *C. pseudodiphtheriticum* was used for the in vitro study, which limits the generalisability of the results. It is possible that other strains of these commensal bacteria may not have the same inhibitory effect on *S. pneumoniae*. Also, the results obtained should be considered in the context of previous findings on the dynamics of bacterial inhibition, where the order of exposure between commensal and pathogenic bacteria plays a crucial role (53). In this study, a bacteria invasion scenario, where a pathogen attempts to colonise a pre-existing respiratory microbiota, was simulated by adding *S. pneumoniae* to a previously grown culture of both commensal bacteria. Future studies could explore more scenarios to determine to what extent the order affects the ability of commensal bacteria to inhibit respiratory pathogens.

Furthermore, a few of the observed associations did not reach statistical significance, which could be explained by the limited sample size. Future research should consider increasing the sample size and incorporating genomic and phenotypic studies to improve the accuracy and elucidate the possible mechanism of action.

These findings could have important implications for understanding microbial interactions within polymicrobial environments, particularly in the context of bacterial interference and competition within the respiratory tract. The observed inhibitory effects suggest that these commensal bacteria may play a protective role in mitigating pneumococcal replication. The antagonistic effect of these commensal bacteria on *S. pneumoniae* replication supports the potential protective factor of a healthy nasopharyngeal microbiota against IPD and underscores the potential of these microorganisms as promising probiotic candidates.

## ACKNOWLEDGES

The authors acknowledge the financial assistance extended by the projects FIS PI19/00104 and PI23/00049, and to PFIS fellowship FI 24/00206. This support was crucial for the attainment of results presented in the present research.

## AUTHOR CONTRIBUTIONS

Conceptualization: C.MA and M.BF; Data curation: M.C; Formal analysis: M.C and M.BF; Investigation: M.C; Methodology: C.MA, P.B, D.H, A.L and A.M; Resources: D.H, A.PA and P.C; Supervision: C.MA and M.BF; Writing-original draft: C.MA, M.C and M.BF; Writing-review and editing: A.L, P.B, D.H, A.PA, P.C, G. GC and A.M.

---

**Table S1:**
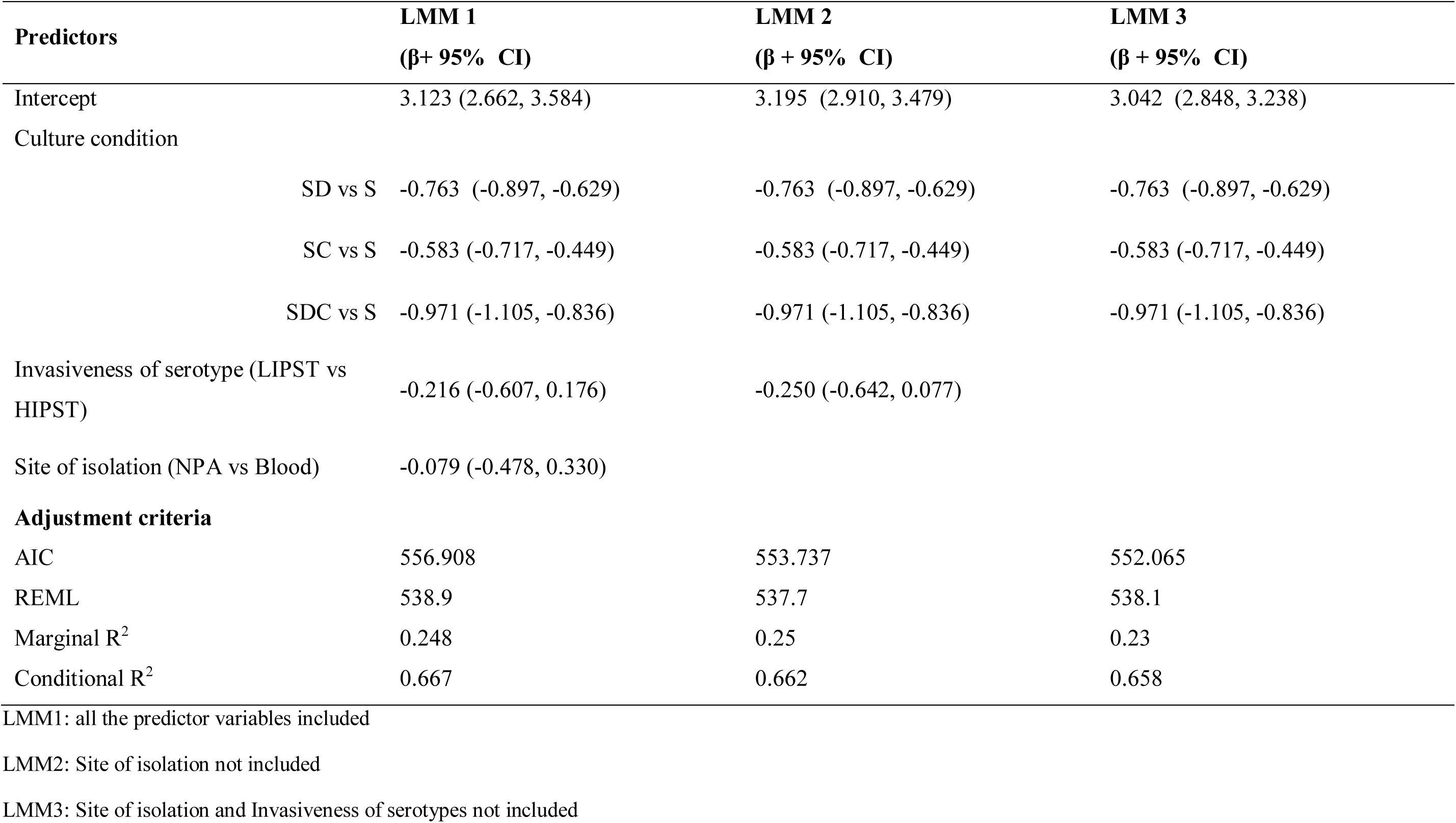
Comparison of the various linear mixed-effect models performed.

**Table S2:** Linear mixed-effect model to analyse the effect of Jnvasiveness of serotype in presence or absence of commensal bacteria to pneumococcal growth.

| Predictors | $\beta$ | 95% CI | SE | p value |
| --- | --- | --- | --- | --- |
| Intercept | 3.111 | 2.806, 3.415 | 0.155 | $p < 0.0001$ **** |
| Culture condition |  |  |  |  |
| SD vs S | -0.662 | -0.877, -0.447 | 0.11 | $p < 0.0001$ **** |
| SC vs S | -0.495 | -0.710, -0.300 | 0.11 | $p < 0.0001$ **** |
| SDC vs S | -0.822 | -1.037, -0.607 | 0.11 | $p < 0.0001$ **** |
| Invasiveness of serotype (LIPST vs HIPST) | -0.112 | -0.642, 0.077 | 0.2 | 0.577 |
| Culture condition*Invasiveness of serotype |  |  |  |  |
| S: LIPST vs HIPST | -0.112 | -0.515, 0.291 | 0.2 | 0.577 |
| SD: LIPST vs HIPST | -0.278 | -0.681, 0.125 | 0.2 | 0.171 |
| SC: LIPST vs HIPST | -0.257 | -0.659, 0.146 | 0.2 | 0.205 |
| SDC: LIPST vs HIPST | -0.355 | -0.758, 0.047 | 0.2 | 0.082 · |
| **Random Effects** |  |  |  |  |
| $\sigma^2$ | 0.197 | | | |
| $\tau_{00}$ Replicate:Strain | 0.063 | | | |
| $\tau_{00}$ Strain | 0.178 | | | |
| ICC | 0.55 |  |  |  |
| $N_{\text{REPLICATE}}$ | 3 | | | |
| $N_{\text{STRAIN}}$ | 28 | | | |
| Observations | 338 |  |  |  |

$\beta$ : Estimated Coefficient
SE: Standard Error
$\sigma^2$ : Residual variance
$\tau_{00}$ : Intercept variance
ICC: Intraclass Correlation Coefficient

## REFERENCES

1. Bogaert D, De Groot R, Hermans PWM. 2004. *Streptococcus pneumoniae* colonisation: the key to pneumococcal disease. Lancet Infect Dis 4:144–154. 10.1016/S1473-3099(04)00938-7

2. Wahl B, O’Brien KL, Greenbaum A, Majumder A, Liu L, Chu Y, Lukšić I, Nair H, McAllister DA, Campbell H, Rudan I, Black R, Knoll MD. 2018. Burden of *Streptococcus pneumoniae* and *Haemophilus influenzae* type b disease in children in the era of conjugate vaccines: global, regional, and national estimates for 2000–15. Lancet Glob Health 6:e744–e757.

3. Morais V, Suarez N. 2022. Conjugation Mechanism for Pneumococcal Glycoconjugate Vaccines: Classic and Emerging Methods. Bioengineering 10.3390/bioengineering9120774.

4. Balsells E, Guillot L, Nair H, Kyaw MH. 2017. Serotype distribution of *Streptococcus pneumoniae* causing invasive disease in children in the post-PCV era: A systematic review and meta-analysis. PLoS One 12. 10.1371/journal.pone.0177113

5. Balsells E, Dagan R, Yildirim I, Gounder PP, Steens A, Muñoz-Almagro C, Mameli C, Kandasamy R, Givon Lavi N, Daprai L, van der Ende A, Trzciński K, Nzenze SA, Meiring S, Foster D, Bulkow LR, Rudolph K, Valero-Rello A, Ducker S, Vestrheim DF, von Gottberg A, Pelton SI, Zuccotti GV, Pollard AJ, Sanders EAM, Campbell H, Madhi SA, Nair H, Kyaw MH. 2018. The relative invasive disease potential of *Streptococcus pneumoniae* among children after PCV introduction: A systematic review and meta-analysis. Journal of Infection 77. 10.1016/j.jinf.2018.06.004

6. Brueggemann AB, Griffiths DT, Meats E, Peto T, Crook DW, Spratt BG. 2003. Clonal relationships between invasive and carriage *Streptococcus pneumoniae* and serotype- and clone-specific differences in invasive disease potential. Journal of Infectious Diseases 187.

7. Weinberger DM, Malley R, Lipsitch M. 2011. Serotype replacement in disease after pneumococcal vaccination. The Lancet 10.1016/S0140-6736(10)62225-8.

8. Hanquet G, Krizova P, Dalby T, Ladhani SN, Nuorti JP, Danis K, Mereckiene J, Knol MJ, Winje BA, Ciruela P, de Miguel S, Portillo ME, MacDonald L, Morfeldt E, Kozakova J, Valentiner-Branth P, Fry NK, Rinta-Kokko H, Varon E, Corcoran M, van der Ende A, Vestrheim DF, Munoz-Almagro C, Sanz JC, Castilla J, Smith A, Henriques-Normark B, Colzani E, Pastore-Celentano L, Savulescu C. 2022. Serotype Replacement after Introduction of 10-Valent and 13-Valent Pneumococcal Conjugate Vaccines in 10 Countries, Europe. Emerg Infect Dis 28.

9. Simell B, Auranen K, Käyhty H, Goldblatt D, Dagan R, O’Brien KL. 2012. The fundamental link between pneumococcal carriage and disease. Expert Rev Vaccines 10.1586/erv.12.53.

10. Laufer AS, Metlay JP, Gent JF, Fennie KP, Kong Y, Pettigrewa MM. 2011. Microbial communities of the upper respiratory tract and otitis media in children. mBio 2. 10.1128/mbio.00245-10

11. Copeland E, Leonard K, Carney R, Kong J, Forer M, Naidoo Y, Oliver BGG, Seymour JR, Woodcock S, Burke CM, Stow NW. 2018. Chronic rhinosinusitis: Potential role of microbial dysbiosis and recommendations for sampling sites. Front Cell Infect Microbiol 8. 10.3389/fcimb.2018.00057

12. Henares D, Brotons P, de Sevilla MF, Fernandez-Lopez A, Hernandez-Bou S, Perez-Argüello A, Mira A, Muñoz-Almagro C, Cabrera-Rubio R. 2021. Differential nasopharyngeal microbiota composition in children according to respiratory health status. Microb Genom 7:661. 10.1099/mgen.0.000661

13. Lappan R, Imbrogno K, Sikazwe C, Anderson D, Mok D, Coates H, Vijayasekaran S, Bumbak P, Blyth CC, Jamieson SE, Peacock CS. 2018. A microbiome case-control study of recurrent acute otitis media identified potentially protective bacterial genera. BMC Microbiol 18.

14. Brugger SD, Eslami SM, Pettigrew MM, Escapa IF, Henke MT, Kong Y, Lemon KP. 2020. *Dolosigranulum pigrum* Cooperation and Competition in Human Nasal Microbiota. mSphere 5. 10.1128/msphere.00852-20

15. Raya Tonetti F, Tomokiyo M, Ortiz Moyano R, Quilodrán-Vega S, Yamamuro H, Kanmani P, Melnikov V, Kurata S, Kitazawa H, Villena J. 2021. The Respiratory Commensal Bacterium *Dolosigranulum pigrum* 040417 Improves the Innate Immune Response to *Streptococcus pneumoniae*. Microorganisms 9. 10.3390/microorganisms9061324

16. Moyano RO, Tonetti FR, Tomokiyo M, Kanmani P, Vizoso-Pinto MG, Kim H, Quilodrán-Vega S, Melnikov V, Alvarez S, Takahashi H, Kurata S, Kitazawa H, Villena J. 2020. The Ability of Respiratory Commensal Bacteria to Beneficially Modulate the Lung Innate Immune Response Is a Strain Dependent Characteristic. Microorganisms 8. 10.3390/microorganisms8050727

17. De Boeck I, Wittouck S, Martens K, Spacova I, Cauwenberghs E, Allonsius CN, Jörissen J, Wuyts S, Van Beeck W, Dillen J, Bron PA, Steelant B, Hellings PW, Vanderveken OM, Lebeer S. 2021. The nasal mutualist *Dolosigranulum pigrum* AMBR11 supports homeostasis via multiple mechanisms. iScience 24. 10.1016/j.isci.2021.102978

18. Islam MA, Albarracin L, Melnikov V, Andrade BGN, Cuadrat RRC, Kitazawa H, Villena J. 2021. *Dolosigranulum pigrum* Modulates Immunity against SARS-CoV-2 in Respiratory Epithelial Cells. Pathogens 10.

19. Stubbendieck RM, Hurst JH, Kelly MS. 2024. *Dolosigranulum pigrum*: A promising nasal probiotic candidate. PLoS Pathog 20:e1011955.

20. Aguirre M, Morrison D, Cookson BD, Gay FW, Collins MD. 1993. Phenotypic and phylogenetic characterization of some Gemella-like organisms from human infections: description of *Dolosigranulum pigrum* gen. nov., sp. nov. J Appl Bacteriol 75:608–612.

21. Popowitch EB, Boiditswe SC, Patel MZ, Aquino JN, Sozat AK, Caiazzo AJ, Maldonado-Barragán A, Hurst JH, Steenhoff AP, Kelly MS. 2024. *Dolosigranulum savutiense* sp. nov., isolated from human upper respiratory samples collected in Botswana. Int J Syst Evol Microbiol 74.

22. LaClaire L, Facklam R. 2000. Antimicrobial Susceptibility and Clinical Sources of *Dolosigranulum pigrum* Cultures. Antimicrob Agents Chemother 44:2001.

23. De Boeck I, Spacova I, Vanderveken OM, Lebeer S. 2021. Lactic acid bacteria as probiotics for the nose? Microb Biotechnol 10.1111/1751-7915.13759.

24. Aziz M, Palmer A, Iversen S, Salazar JE, Pham T, Roach K, Becker K, Kaspar U, Price LB, Baig S, Stegger M, Andersen PS, Liu CM. 2023. Design and validation of *Dolosigranulum pigrum* specific PCR primers using the bacterial core genome. Sci Rep 13.

25. Flores Ramos S, Brugger SD, Escapa IF, Skeete CA, Cotton SL, Eslami SM, Gao W, Bomar L, Tran TH, Jones DS, Minot S, Roberts RJ, Johnston CD, Lemon KP. 2021. Genomic Stability and Genetic Defense Systems in *Dolosigranulum pigrum*, a Candidate Beneficial Bacterium from the Human Microbiome. mSystems 6.

26. Mokoena MP. 2017. Lactic acid bacteria and their bacteriocins: Classification, biosynthesis and applications against uropathogens: A mini-review. Molecules 10.3390/molecules22081255.

27. Yildiz S, Bonifacio Lopes JPP, Bergé M, González-Ruiz V, Baud D, Kloehn J, Boal-Carvalho I, Schaeren OP, Schotsaert M, Hathaway LJ, Rudaz S, Viollier PH, Hapfelmeier S, Francois P, Schmolke M. 2020. Respiratory tissue-associated commensal bacteria offer therapeutic potential against pneumococcal colonization. Elife 9.

28. Burkovski A. 2015. Corynebacterium pseudodiphtheriticum: Putative probiotic, opportunistic infector, emerging pathogen. Virulence 10.1080/21505594.2015.1067747.

29. Singhal N, Kumar M, Kanaujia PK, Virdi JS. 2015. MALDI-TOF mass spectrometry: An emerging technology for microbial identification and diagnosis. Front Microbiol 10.3389/fmicb.2015.00791.

30. Centers for Disease Control and Prevention. 2011. Primary Culture and Presumptive Identification of Neisseria meningitidis, *Streptococcus pneumoniae*, and *Haemophilus influenzae*. Laboratory Methods for the Diagnosis of Meningitis.

31. Selva L, Del Amo E, Brotons P, Muñoz-Almagro C. 2012. Rapid and easy identification of capsular serotypes of *Streptococcus pneumoniae* by use of fragment analysis by automated fluorescence-based capillary electrophoresis. J Clin Microbiol 50.

32. Liyanapathirana V, Ang I, Tsang D, Fung K, Ng TK, Zhou H, Ip M. 2014. Application of a target enrichment-based next-generation sequencing protocol for identification and sequence-based prediction of pneumococcal serotypes. BMC Microbiol 14.

33. Carvalho MDGS, Tondella ML, McCaustland K, Weidlich L, McGee L, Mayer LW, Steigerwalt A, Whaley M, Facklam RR, Fields B, Carlone G, Ades EW, Dagan R, Sampson JS. 2007. Evaluation and improvement of real-time PCR assays targeting lytA, ply, and psaA genes for detection of pneumococcal DNA. J Clin Microbiol 45.

34. Santoro F, Iannelli F, Pozzi G. 2019. Genomics and Genetics of *Streptococcus pneumoniae*. Microbiol Spectr 7.

35. Portet S. 2020. A primer on model selection using the Akaike Information Criterion. Infect Dis Model 5.

36. Lin B, Pang Z, Jiang J. 2013. Fixed and random effects selection by REML and pathwise coordinate optimization. Journal of Computational and Graphical Statistics 22.

37. Edwards LJ, Muller KE, Wolfinger RD, Qaqish BF, Schabenberger O. 2008. An R2 statistic for fixed effects in the linear mixed model. Stat Med 27.

38. Kim JH. 2019. Multicollinearity and misleading statistical results. Korean J Anesthesiol 72.

39. R Core Team. 2019. R: A language and environment for statistical computing. R Foundation for Statistical Computing.

40. Weisberg S, Fox J. 2011. An R Companion to Applied RegressionSage.

41. Bates D, Mächler M, Bolker BM, Walker SC. 2015. Fitting linear mixed-effects models using lme4. J Stat Softw 67.

42. Lenth R V. 2024. emmeans: Estimated Marginal Means, aka Least-Squares Means. R package version 1102090002.

43. Kuznetsova A, Brockhoff PB, Christensen RHB. 2017. lmerTest Package: Tests in Linear Mixed Effects Models. J Stat Softw 82.

44. Bartoń K. 2009. MuMIn: Multi-model inference. R package version 1436.

45. Belman S, Lefrancq N, Nzenze S, Downs S, du Plessis M, Lo SW, McGee L, Madhi SA, von Gottberg A, Bentley SD, Salje H, Corso A, Gagetti P, Brooks AW, Hasanuzzaman M, Saha SK, Saha S, Davydov A, Titov L, Almeida SCG, Turner P, Zhao C, Wang H, Ip M, Ho PL, Law P, Keenan JD, Cohen R, Varon E, Sampane-Donkor E, Veeraraghavan B, Nagaraj G, Ravikumar KL, Yuvaraj J, Shamanna Noga V, Benisty R, Dagan R, Bigogo G, Verani J, Kiran A, Everett DB, Cornick J, Alaerts M, Sekaran SD, Clarke SC, Moiane B, Sigauque B, Mucavele H, Pollard AJ, Kandasamy R, Carter PE, Obaro SK, Lehmann D, Ford R, Ochoa TJ, Skoczynska A, Sadowy E, Hryniewicz W, Puzia W, Doiphode S, Egorova E, Voropaeva E, Urban Y, Kastrin T, Ndlangisa K, De Gouveia L, Ali M, Wolter N, Lekhuleni C, Almagro CM, Alonso AR, Henares D, Srifuengfung S, Kwambana-Adams B, Foster-Nyarko E, Bojang E, Antonio M, Tientcheu PE, Moïsi J, Nurse-Lucas M, Akpaka PE, Eser ÖK, Scott A, Aanensen D, Croucher N, Lees JA, Gladstone RA, Tonkin-Hill G, Chaguza C, Cleary D, Mellor K, Beall B, Klugman KP, Rodgers G, Hawkins PA, Blaschke AJ, Pershing NL. 2024. Geographical migration and fitness dynamics of *Streptococcus pneumoniae*. Nature 631:386–392.

46. Stubbendieck RM, May DS, Chevrette MG, Temkin MI, Wendt-Pienkowski E, Cagnazzo J, Carlson CM, Gern JE, Currie CR. 2019. Competition among nasal bacteria suggests a role for siderophore-mediated interactions in shaping the human nasal microbiota. Appl Environ Microbiol 85.

47. Hernández-González JC, Martínez-Tapia A, Lazcano-Hernández G, García-Pérez BE, Castrejón-Jiménez NS. 2021. Bacteriocins from lactic acid bacteria. A powerful alternative as antimicrobials, probiotics, and immunomodulators in veterinary medicine. Animals 10.3390/ani11040979.

48. Meade E, Slattery MA, Garvey M. 2020. Bacteriocins, potent antimicrobial peptides and the fight against multi drug resistant species: Resistance is futile? Antibiotics 10.3390/antibiotics9010032.

49. Schofs L, Sparo MD, Sánchez Bruni SF. 2020. Gram-positive bacteriocins: usage as antimicrobial agents in veterinary medicine. Vet Res Commun 10.1007/s11259-020-09776-x.

50. Sanchez-Rosario Y, Johnson MDL. 2021. Media Matters, Examining Historical and Modern *Streptococcus pneumoniae* Growth Media and the Experiments They Affect. Front Cell Infect Microbiol 10.3389/fcimb.2021.613623.

51. Renz A, Widerspick L, Dräger A. 2021. First genome-scale metabolic model of *Dolosigranulum pigrum* confirms multiple auxotrophies. Metabolites 11.

52. Tran TH, Escapa IF, Roberts AQ, Gao W, Obawemimo AC, Segre JA, Kong HH, Conlan S, Kelly MS, Lemon KP. 2023. Metabolic capabilities are highly conserved among human nasal-associated *Corynebacterium* species in pangenomic analyses 10.1101/2023.06.05.543719.

53. Díaz-Garrido N, Lozano CP, Kreth J, Giacaman RA. 2020. Competition and Caries on Enamel of a Dual-Species Biofilm Model with *Streptococcus mutans* and *Streptococcus sanguinis*. Appl Environ Microbiol 86.

